# Respiratory strategy at birth initiates distinct lung injury phenotypes in the preterm lamb lung

**DOI:** 10.1101/2022.09.06.506865

**Authors:** Prue M. Pereira-Fantini, Kristin Ferguson, Karen McCall, Regina Oakley, Elizabeth Perkins, Sean Byars, Nicholas Williamson, Shuai Nie, David G Tingay

## Abstract

Bronchopulmonary disease is the chronic manifestation of the acute injury that may accompany ventilation following preterm birth. A lack of clear trial evidence often hampers clinical decision-making during support of the preterm lung at birth. Protein biomarkers have been used to define acute lung injury phenotypes and improve patient selection for specific interventions in adult respiratory distress syndrome. Here we present a mass spectrometry-based approach to profile the protein phenotype associated with three different aeration strategies known to cause different pathophysiological responses when applied at birth to preterm lambs. We were able to identify pathway enrichments specific to both ventilation strategy and lung regions based upon gravity-dependency. Ventilation strategy-specific phenotypes were further delineated by applying partial least square modelling to identify associations between specific proteins and clinical, physiological and morphological outcomes. This work highlights the specificity of lung injury responses to routinely applied birth interventions such as different respiratory support approaches and identified the molecular events associated with each. Furthermore, we demonstrate the capacity to subdivide preterm infants by the direct aetiology and response to lung injury; the first step towards true precision medicine in neonatology.

## INTRODUCTION

Bronchopulmonary dysplasia (BPD) remains a serious chronic complication of preterm birth, with potentially life-long consequences. BPD arises from acute injurious events occurring in the under-developed preterm lung, often after ventilation begins at birth (1, 2). This includes inappropriate ventilatory choices aimed at supporting the respiratory distress syndrome (RDS) characteristic of preterm birth (3). The complex and dynamic cascade of injury is unlikely to be explained by a singular biological or mechanical cause. Yet, trials of preterm respiratory therapies are generally premised on a singular, unified treatment model, and have generally failed to show any impact on BPD rates (4, 5). The characterisation of acute lung injury and BPD is also often reliant on imprecise clinical and diagnostic criteria, such as oxygen dependency (6). This approach fails to appreciate the complex interaction between intrinsic and developmental factors and clinical interventions that shapes the development and progression from acute lung injury to BPD, nor does it enable a biological understanding of the individual patients response to treatment.

A possible solution has been explored in adults with acute respiratory distress syndrome (ARDS); using physiology, clinical data, biomarkers to stratify patients into distinct, homogenous subgroups (7). This approach has identified ARDS phenotypes associated with diverging inflammatory responses (8–10) that indicate different benefit response to specific therapies, such as lung recruitment (11). Considering ARDS phenotypes in this way provides insight into the variable outcomes from ARDS trials, and allows the potential for targeted interventions (8). Like ARDS, the preterm RDS population exhibits a high degree of clinical and biological heterogeneity, suggesting that this population may similarly benefit from identification of biomarker-based phenotypes.

Effectively achieving lung aeration at birth is fundamental to establishing a functional residual capacity, and failing to so do correlates to the earliest point in which respiratory management can initiate lung injury (12, 13). Various approaches that support aeration in preterm infants have been suggested targeting different physiological rationales, such as positive end-expiratory pressure (PEEP) strategies, or the use of an initial sustained lung inflation (SI) (12, 14–16). It has been proposed that the different clinical and injury responses seen with these approaches reflect different injury mechanistic pathways within the lung, and thus differing phenotypes which are as yet undefined.

We have established a large biobank of lung tissue, clinical and lung imaging and mechanics data in preterm lambs. We have also previously identified protein-based phenotypes associated with gestational age (17) and duration of ventilation (18) in this biobank. Thus, the primary aim of this study was to use proteomics to provide a deeper biological understanding of acute lung injury phenotypes resulting from different aeration strategies at birth in the preterm lung with RDS. This aim will be achieved by comparing the phenotypic response to different PEEP strategies at birth in the preterm lamb. In secondary studies, we aimed to use partial least squares modelling to identify phenotype specific protein-function associations within non-dependent and dependent lung regions. By doing so we aim to design the first phenotypic approach to defining the initiation of preterm lung disease. This would provide the mechanistic foundation for refining lung protective decisions following preterm birth to the most appropriate patient or intervention-based group, and potentially provide the precision needed to meaningfully reduce the incidence of BPD (3).

## RESULTS

Lambs were well-matched clinically (Supplemental Table 1).

### Proteome profiling identified regional and ventilation strategy-specific alterations in the lung proteome

2373 proteins were analysed. The total number of differentially expressed proteins (DEPs) within the lung was highest in the SI group, followed by No-RM (Figure 1A). The non-gravity dependent lung had more DEPs than the dependent lung for all ventilation groups. Co-expression of DEPs in both lung regions was approximately 10-fold higher in No-RM and SI groups compared to DynPEEP (Figure 1B). 56% (100 proteins) of DEPs in the non-dependent lung and 69% (14 proteins) of DEPs in the dependent lung were only expressed in a single study group (Figure 1C). There were a higher number of decreased DEPs than increased DEPs in both lung regions of the SI and DynPEEP groups and the opposite for No-RM group (Figure 1D). A total of 20 protein classes were assigned to the non-dependent lung and 18 the dependent lung (Figure 1E). In the non-dependent lung 15 protein classes were common to all intervention groups (Supplemental Table 2), with defence/immunity proteins only identified in the SI group and cell adhesion molecules, structural proteins and transmembrane signal receptors absent in the DynPEEP group. Translational proteins were proportionally two-fold higher in the DynPEEP group compared to the No-RM and SI groups (23% *versus* 11 and 12%). In the dependent lung, protein class diversity was decreased by approximately 45% in the DynPEEP group compared to the other groups, except for transporter (3-fold increase) and gene-specific transcriptional regulator proteins (3 to 6-fold)(Supplemental Table 3).

**Figure 1:**
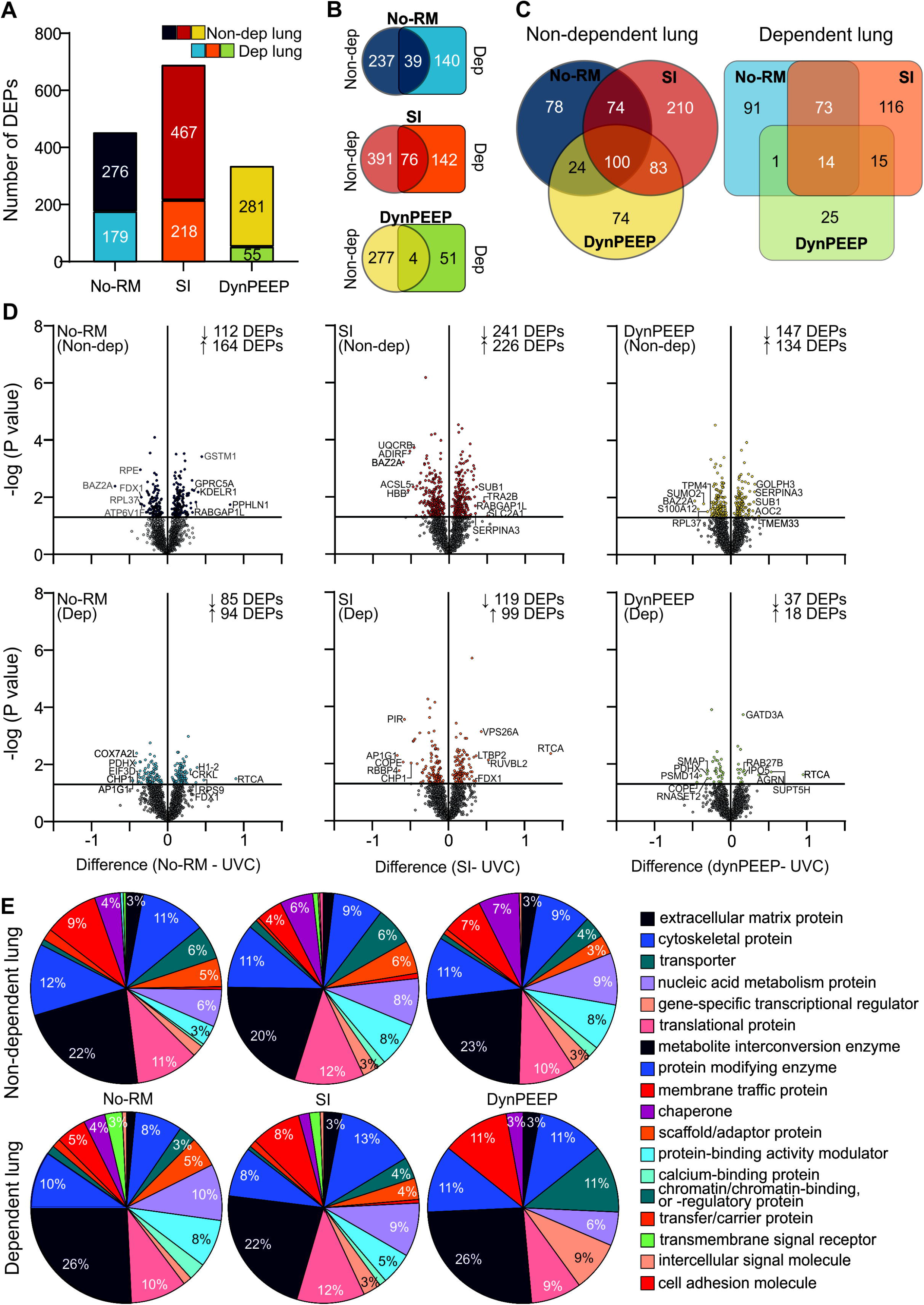
Results from Orbitrap-MS analysis at 90 minutes of life in the non-dependent and dependent lung for the no-recruitment manoeuvre (No-RM), sustained inflation (SI) and dynamic stepwise positive end-expiratory pressure (DynPEEP) strategies relative to unventilated controls. Number of differentially expressed proteins (DEPs) (A), Venn diagram comparing differential protein expression between lung regions (B) and between intervention strategies within the same lung region (C). Volcano plots with DEPs highlighted in colour and the five most abundance and depleted proteins identified (D). PANTHER analysis of the protein class composition amongst DEPs identified in non-dependent and dependent lung (E).

### WebGestalt over-representation analysis (ORA) revealed lung region specific pathway enrichment, with unique pathways associated with specific ventilation strategies

In cellular enrichment analysis, non-dependent lung DEPs from the DynPEEP group were cytoplasmic (Figure 2A). Uniquely, only DEPs from the No-RM group were associated with adherens and cell-substrate junction, and only SI DEPs were from the extracellular region. Both the No-RM group and SI group were associated with the cell membrane and organelle components. A broader array of cellular components was identified in the dependent lung, largely due to association with specific organelle components in the No-RM (mitochondria, z disc), SI (endoplasmic reticulum, golgi and actin cytoskleleton) and DynPEEP (organelle membrane and secretory vesicle) groups.

**Figure 2:**
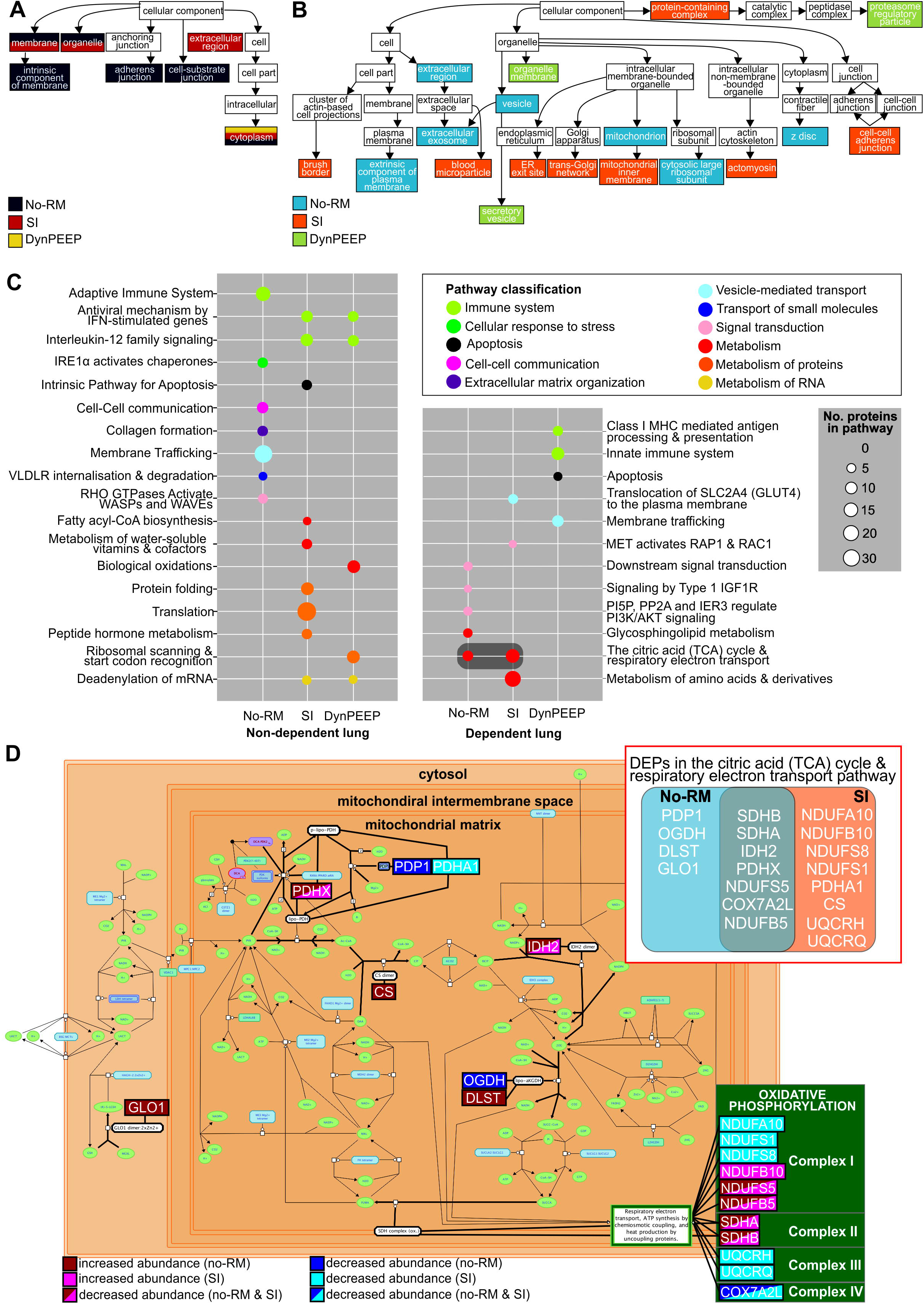
WebGestalt identification of enriched cellular components and biological pathways in non-dependent and dependent lung exposed to either the no-recruitment manoeuvre (No-RM), sustained inflation (SI) or dynamic stepwise positive end-expiratory pressure (DynPEEP) strategy. Enrichment of cellular components in the nondependent and dependent lung (A). Identified Reactome pathways with dark grey box highlighting co-enrichment of *the citric acid (TCA) cycle and respiratory electron transport pathway* in both No-RM and SI dependent lung (B). DEP mapping of No-RM and SI DEPs onto *the citric acid (TCA) cycle and respiratory electron transport pathways* (C). Inset; Venn diagram comparing DEP identification within the pathway in No-RM and SI groups. The threshold for identification of significant enrichments was adjusted p value < 0.05 and three or more pathways associated with the cellular component or pathway. Redundant Reactome pathways were reduced using weighted set cover. High resolution version of Panel A and C available in the online supplement.

18 enriched Reactome pathways were identified in the non-dependent lung and 12 in the dependent lung (Figure 2B). In the non-dependent lung, the inflammation response was divergent amongst RM-groups with the *adaptive immune system* enriched only in the No-RM group, as opposed to *cytokine signalling pathway* enrichments in SI and DynPEEP groups. Several pathways were enriched only in a single RM group and these included; *Membrane trafficking* and *collagen formation* (No-RM), *Apoptosis* and *Translation* (SI) and *Biological oxidations* and *Ribosomal scanning* (DynPEEP).

In the dependent lung, enriched pathways were specific to RM-groups with the exclusion of *the citric acid cycle (TCA) and respiratory electron transport pathway*, which was enriched in both the No-RM and SI groups. Amongst the DEPs associated with *The TCA pathway*, seven DEPs were altered in both the No-RM and SI groups, four DEPs were altered in only the No-RM group and eight DEPs were altered in only the SI group. 58% of DEPs mapped to the oxidative phosphorylation component of the *TCA and respiratory electron transport pathway* (Figure 2C), with No-RM DEPs associated with Complexes I and II, whilst SI DEPs were associated with Complexes I-IV.

### Integrative analysis identified regional and temporal protein-function associations specific to each ventilation strategy

In addition to comparing the fold change in protein abundance between ventilated and unventilated lung, we also used sPLS to test for associations between protein abundance (normalised protein reporter intensities reported in the ventilated lung) and functional measures of aeration, ventilation and clinical status. Two protein-function networks were observed in all lung regions and ventilation strategies, with an additional single protein-function network observed in the non-dependent lung of the DynPEEP group (Table 1). The highest number of protein-function associations were observed in the No-RM group (Table 1).

**Table 1:** Summary of regional and temporal outcomes from sparse partial least squared correlations (sPLS) between protein abundance and functional measures. A cutoff threshold of 0.7 (positive and negative) was applied to the analysis.

|  | No-RM |  | SI |  | DynPEEP |  |
| --- | --- | --- | --- | --- | --- | --- |
|  | Non-dep | Dep | Non-dep | Dep | Non-dep | Dep |
| No. networks | 2 | 2 | 2 | 2 | 3 | 2 |
| No. proteins | 20 | 20 | 16 | 4 | 8 | 12 |
| No. measures | 19 | 15 | 8 | 9 | 9 | 5 |
| Temporal functional observations |  |  |  |  |  |  |
| 3 minutes (proportion; %) | 3 (16) | 0 (0) | 1 (13) | 3 (33) | 3 (33) | 0 (0) |
| 5 minutes (proportion; %) | 6 (32) | 2 (13) | 6 (75) | 3 (33) | 3 (33) | 1 (20) |
| 90 minutes (proportion; %) | 10 (53) | 13 (87) | 1 (13) | 3 (33) | 3 (33) | 4 (80) |

Distinct temporal associations were observed between lung regions and ventilation strategies (Table 1, Figure 3). In the No-RM group the highest number of protein-function associations in either the non-dependent or dependent lung were observed after 90 minutes of respiratory support, and included measures of lung morphology, lung injury and oxygenation and acid-base. In contrast, in the SI group, ventilation parameters and C_dyn_ at 5 minutes accounted for 71% of protein-function relations in the non-dependent lung. In the dependent lung a single protein network was associated with the completion of the SI and 5 minute ventilation parameters and C_dyn_ and a second network consisting of three proteins associated with acid-base at 90 minutes. The DynPEEP group also exhibited associations between non-dependent lung proteins and ventilation parameters at the end of the DynPEEP strategy and 5 minutes, as well as 90 minutes. 80% of associations in the DynPEEP dependent lung involved functional measures at 90 min (oxygenation and haemoglobin).

**Figure 3:**
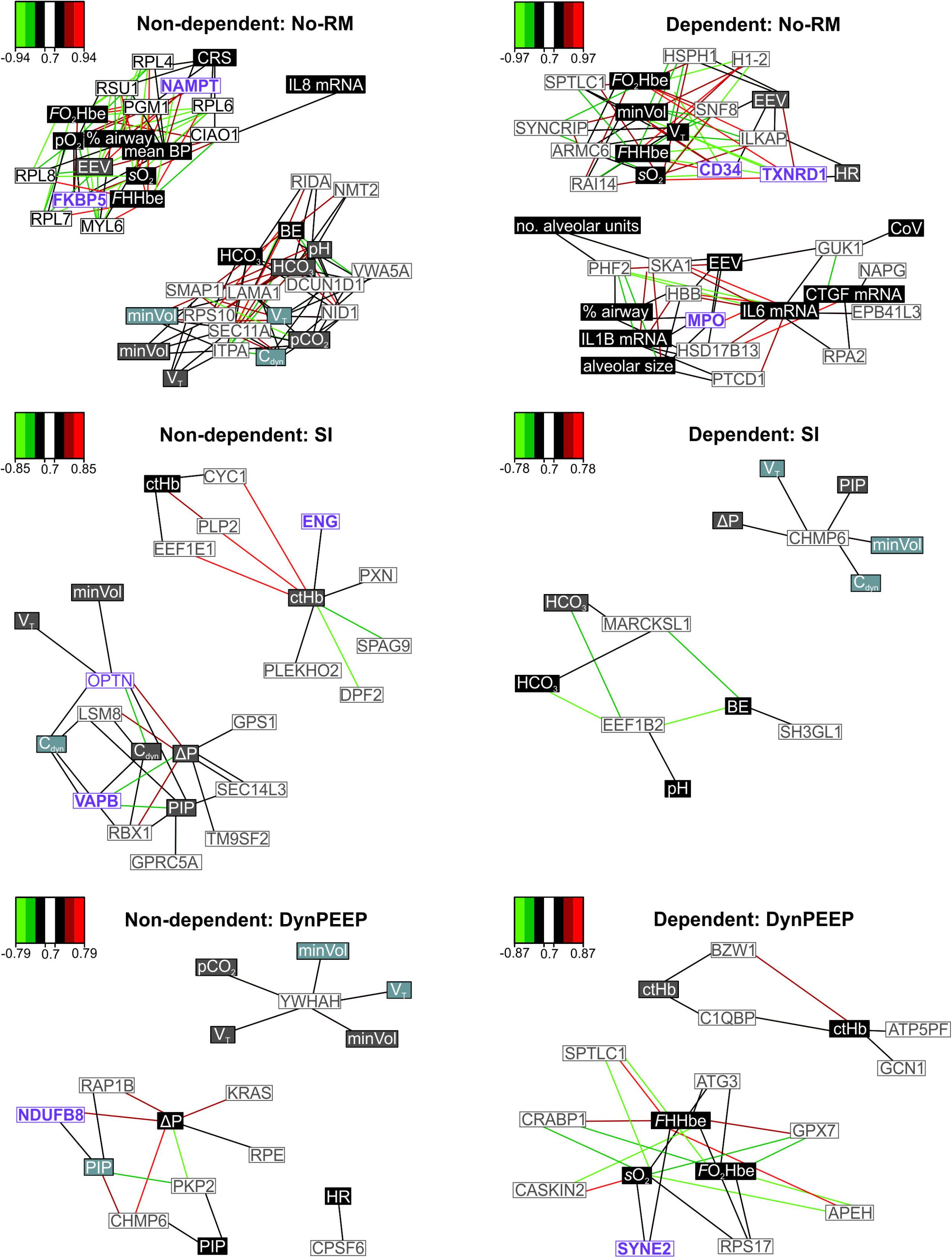
Spared partial least squared correlations (sPLS) between protein abundance and functional measures in the non-dependent and dependent lung of ventilated preterm lambs receiving no-recruitment manoeuvre (No-RM), sustained inflation (SI) or a dynamic stepwise positive end-expiratory pressure (DynPEEP) strategy. sPLS in regression model was used to model a causal relationship between the proteins and functional measures. The network is displayed graphically as nodes (proteins and functional measures) and edges (relationship between nodes), with the colour edge intensity indicating the level of the association: red, positive and green, negative. Only the strongest pairwise associations were displayed, with a cut-off threshold of 0.7 (positive and negative). Coloured boxes indicate timing of functional measures (light blue = during recruitment manoeuvre (first 3 minutes), grey = 5 minutes post-birth, black = 90 minutes post-birth). White boxes contain the protein name with proteins identified in the DisGeNET database as associated with bronchopulmonary dysplasia, respiratory distress syndrome or respiratory failure highlighted in bold purple text. *Function measure abbreviations; C_RS_, total compliance; BE, base excess; V_min_, minute volume; V_T_, tidal volume; C_dyn_, dynamic compliance; PIP, peak inspiratory pressure; ΔP, change in pressure; HR, heart rate; mean BP, mean blood pressure*.

## DISCUSSION

To our knowledge this is the first study to map the complete protein-functional responses of the preterm lung at initiation of aeration and ventilation at birth, the earliest time point in which clinical respiratory decisions can impact injury. Effective establishment of ventilation at birth is vital (30). The optimal approach to support ventilation in the preterm lung whilst minimising injury remains unclear due to crude, late and in-direct measures of lung injury, such as death and oxygen requirement. These fail to delineate the molecular impact of such strategies on the early, acute lung injury response (31). We have recently demonstrated that lung mechanics, regional aeration and ventilation states (such as relative overdistension and atelectasis), as well as gravity dependent expression of early injury markers, differed between SI, No-RM and DynPEEP strategies, suggesting each generates a different, and biologically important, injury response in the preterm lung (12, 19, 23). Interestingly oxygenation, the primary clinical tool for defining clinical status and efficacy in trials, was not a delineator of the injury differences. Thus, our biobank was ideal to demonstrate the role of proteomic analysis to understand the response of the preterm lung to respiratory support before developing clinical trials, and use translational models to better understand the phenotype responses to different clinical approaches to the initial management of RDS.

Overall, the biggest impact on the proteome was seen in the SI group which exhibited the highest number of DEPs, whilst the DynPEEP group had fewer DEPs and a simpler protein class composition. Further highlighting the influence of ventilation strategies on protein expression, only 16% of DEPs in the non-dependent lung and 4% in the dependent lung were common to all aeration strategy groups. Protein expression exhibited a gravity-dependent pattern in all ventilation groups, with more DEPs identified in the gravity non-dependent lung than dependent, and at the individual protein level more than 88% of DEPs were identified in a single lung region alone. We have previously shown that an SI generated the greatest gravity-dependent heterogeneity in aeration and ventilation at birth that caused lung injury events specific to the regional volume state (12). Thus, providing a mechanistic link to explain the greater DEPs especially in the non-dependent lung following a SI. In low compliance states (such as RDS) the non-dependent lung is preferentially ventilated, creating high volutrauma risk during the large mechanical forces applied during the rapid inflation of a SI. A recent large trial of SI at birth was ceased early due to a higher early mortality rate in the SI group of unknown causation (15). Our findings, combined with the previous adverse impacts on regional aeration and surfactant efficacy in preclinical studies (12, 23, 32), provide the first indication that a SI may generate unique and acute biologically harmful changes in the preterm lung related to heterogenous rapid pressure and volume exposure throughout the lung.

Distinct gravity-dependent pathway over-representation was identified for each of the aeration strategies, suggesting differing biological events. In the non-dependent lung there were no common pathways identified between the No-RM and other groups, and only three common pathways were identified for SI and DynPEEP groups. In the dependent lung, only a single common pathway (R-HSA-1428517; citric acid cycle and respiratory electron transport) was identified in the No-RM and SI group. In the non-dependent region of the No-RM lung alone we observed unique identification of 30 proteins associated with an adaptive immune response phenotype. This diverse inflammatory phenotype response after only a brief period of respiratory support may provide insight into why broad anti-inflammatory agents such as corticosteroids, rather than targeted anti-inflammatory therapies, have been effective in reducing the respiratory burden of preterm RDS (33).

The SI non-dependent lung phenotype was strongly and uniquely associated with enrichment of translational proteins with 75% (15/20) of decreased proteins identified as ribosomal proteins and 67% (10/18) of increased proteins as translational proteins (initiation, elongation and release factors). Ribosome biogenesis and protein translation are finely coordinated with and essential for cell growth, proliferation, differentiation, and development (34). Impairment of any of these two cellular processes can severely retard cell growth and perturb development (34). Only 12 proteins were associated with a single pathway in either non-dependent (biological oxidations) or dependent lung (innate immune system) following a DynPEEP strategy. This concurs with our previous observation of improved ventilation and aeration homogeneity, lung mechanics, surfactant efficacy and subsequently reduced early lung injury mRNA markers following DynPEEP compared to SI and No-RM in our biobank (12, 32). Further it provides potential mechanistic support to our hypothesis that gentle intermittent lung aeration via tidal inflations, using PEEP to maintain aeration-gains, will be less injurious than rapid lung inflation (12).

In the dependent lung the TCA cycle and respiratory electron transport pathway was identified as significant in both No-RM and SI dependent lung tissue, suggesting that these aeration strategies may generate significant oxidative stresses. Newborn infants encounter a dramatic surge of oxidative stress immediately after birth. The preterm lung lacks antioxidant protection (35), the risk is further exacerbated when higher oxygen concentrations are used (36). Despite oxidative stress clearly playing a vital role in causing lung injury, antioxidant treatment has not shown clinical efficacy in preventing BPD, suggesting more complex mechanisms are involved (36, 37). In our study we identified nuanced differences in protein abundance profiles between No-RM and SI lung tissues such that whilst complexes II and IV proteins exhibited similar alterations in both strategies, only SI caused decreased abundance of complex I and III proteins. Decreased enzymatic activity of complex I has previously been associated with severity of lung-stretching in a ventilated neonatal mouse model. Conversely, gentler tidal volumes abrogate mitochondrial dysfunction (38). In the current context this suggests that therapies targeting complex I maybe more suited to strategies prone to over-distension such as SI.

In secondary studies we used sparse partial least squares modelling to identify the most important function-protein associations between 61 functional variables from four time-points and protein abundance in 2,373 gravity non-dependent and dependent lung protein. There are several advantages to using this approach; dimension reduction and variable selection is performed simultaneously and allows for identification of most relevant associations, sPLS regression exhibits good performance even when the sample size is much smaller than the total number of variables (as occurs with large ‘omic datasets) and both univariate and multivariate responses can be included in the analysis (39). Highly complex and dense protein-function networks were observed in both lung regions of the No-RM group with most implicated functional measures involving clinical outcome measures and gene expression of lung injury. In contrast both the SI and DynPEEP groups were simpler, with protein-function associations and increased focus on associations between protein abundance and early ventilation delivery. This at first is a surprising result given that the proteome of SI lung tissue was associated with the highest numbers of DEPs. However whilst proteomic analysis facilitated the identification of proteins not previously associated with preterm lung injury, previous preterm, lung injury studies have made a priori assumptions about functional variables, such as the utility of the commonly used 6-member lung injury gene panel, first validated in a No-RM model 12 years ago (40). Several recent papers have used plasma biomarker panels to define broad sub-phenotypes within ARDS (9, 41, 42). Plasma protein biomarkers in general contributed more prominently to the phenotype definitions than most of the commonly used clinical variables in adults with ARDS (9). Few delivery room studies have included protein biomarkers in their assessment, and when they have, single targets were used (43, 44), thereby risking missing unrecognised differences between interventions, and limiting understanding of the pathophysiology events. The diversity in protein phenotype between the three aeration strategies investigated suggests that using non-targeted approaches such as proteomics in early preterm life studies would be highly advantageous at capturing the biological impact of diverse respiratory support strategies.

### Limitations

There are several limitations of this study. We, and others, have previously detailed the clinical translation limitations of our ventilation strategies and instrumented lambs generally (12, 19, 20, 23), specifically suppression of breathing and intubation (19, 22, 23, 45–48). Lamb numbers were unbalanced between the strategies as fewer No-RM study animals, which acted as a ventilation ‘control’ group across studies, were available within the biobank (12, 19, 20, 23). To avoid selection bias and knowing some samples would be lost during the extraction workflow, we included all No-RM, SI and DynPEEP samples rather than matching group sizes. We did not apply FDR correction on the protein p values, however we attempted to counter this within the bioinformatic analysis by limiting reporting of ORA results to cellular compartments and pathways containing at least three DEPs. The sPLS approach being based on protein abundance rather than differential expression does not require consideration of multiple testing principles. To facilitate interpretation of protein-function networks we used the sPLS approach and limited the number of variables to be included and applied an r cutoff of 0.7. Whilst this provided a clearer picture of the variables most contributing to each phenotype, important but weaker biological associations may have been missed.

### Conclusion

Ideally, strategies for BPD prevention should start immediately after birth (49), focusing on supporting lung aeration and then managing RDS whilst minimising ventilator induced lung injury (4). This study is the first to demonstrate that proteomics can identify distinct acute injury responses to different respiratory strategies at birth. Importantly, it maybe possible to use proteomics in the first hours of life to phenotype preterm infants based on the direct aetiology of their lung injury risk, an important step towards true precision respiratory support.

## METHODS

This study used historic samples and data from our biobank. The measurements, physiological data acquisition and analysis and lung injury analysis methodology from these studies have been described previously (12, 19, 20). Figure 4 provides a summary of the experimental workflow and setup relevant to the study aims. All data used in this study is available at Figshare (https://doi.org/10.26188/19252103.v1).

**Figure 4.**
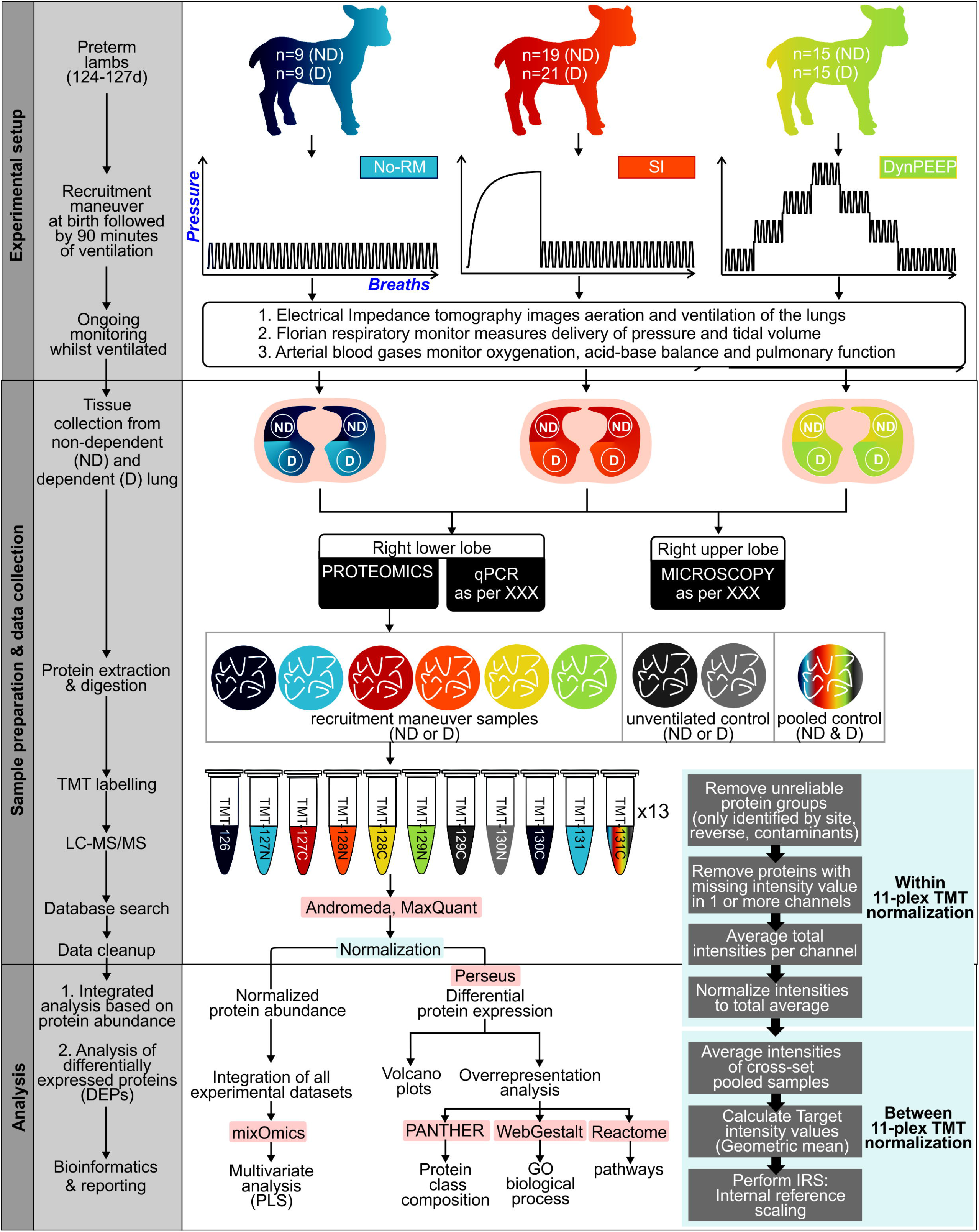
Experimental setup and workflow for comparison of RM-phenotypes in non-dependent (ND) and dependent (D) lung tissue (Main box). The two-step normalization workflow (50) was performed prior to data analysis (Inset box). Blue shading represents No-RM group, red-orange SI group and yellow-green DynPEEP. Software analysis packages highlighted in pink.

### Experimental instrumentation

Preterm lambs (124-127d; term ~145d) born by caesarean section to date-mated, betamethasone-treated ewes were instrumented before delivery (12, 19, 20).

### Recruitment strategies

Before delivery, lambs were randomly assigned to receive one of the following lung recruitment strategies from birth.

#### No intentional recruitment manoeuvre (No-RM)

Positive-pressure ventilation (PPV; SLE5000, SLE Ltd, South Croydon, UK) in volume-targeted ventilation (VTV) mode (PEEP 8 cmH_2_O (21), inspiratory time 0.4 s, rate 60 inflations/min, set V_T_ 7 mL/kg and maximum peak inspiratory pressure (PIP) 35 cmH_2_O).

#### Sustained Inflation (SI)

SI was maintained at 35-40 cmH_2_O until 10 s after achievement of a volume plateau on the EIT display (maximum 180 s) (12, 19, 20, 22).

#### Dynamic PEEP (DynPEEP)

Commencing at 6 cmH_2_O during PPV+VTV (V_T_, 7 ml/kg; maximum PIP 35 cmH_2_O), PEEP increased by 2 cmH_2_O every 10–15 inflations during PPV to a maximum PEEP (14–20 cmH_2_O) based upon dynamic compliance (C_dyn_) response, and then similarly decreased to a final PEEP of 8 cmH_2_O (170–180 s total duration) (12, 19).

On completion of the SI or DynPEEP, PPV+VTV was continued as per the No-RM group for 90 min (12, 19, 20). All lambs commenced PPV using an inspired oxygen fraction (FiO_2_) of 0.3. FiO_2_ and delivered V_T_ were adjusted to maintain peripheral oxygen saturation between 88–94% and a partial pressure of carbon dioxide (PaCO_2_) of 40–60 mmHg using a standardized strategy (12, 19, 23). Poractant alfa 200 mg/kg (Chiesi Farmaceutici SpA) was administered at 10 min. At the end of the study period all lambs received a lethal dose of pentobarbitone (12, 19, 20, 22). Gestation-matched unventilated control (UVC) lambs received a lethal dose of pentobarbitone at delivery for identification of ventilation-associated protein alterations. Immediately after animals were euthanized, the lungs were removed en bloc. Lung tissue samples for proteome analysis from the gravity-dependent and non-dependent zones of the right upper lobe were snap frozen and stored at –80°C until analysis.

### Determination of protein composition in TMT-labelled lung tissues

A detailed methodology describing protein extraction, TMT labelling and data extraction of the preterm lung tissue is available (24). Inclusion in the study was based upon lung samples from our biobank that met the following criteria; ventilation strategy (No-RM, SI or DynPEEP), gestational age <128d, no evidence of illness in ewe (hypoxia, sepsis, chorioamnionitis) and complete qPCR dataset. Following protein extraction only samples with a peptide quantity >10μg were labelled with TMT. Details of these sample reductions and final sample sizes for analysis are shown in Supplementary Figure 1. This provided a final sample of 6-7 lambs in the unventilated control group and between 9-22 in the ventilated groups. As the first study to compare different respiratory interventions in the preterm lung using this methodology a formal power calculation was not possible. We have previously demonstrated that a sample size of 7 unventilated controls and 10 ventilated lambs identified important differences between unventilated and ventilated lambs (18, 25).

### Bioinformatics and statistical analysis

#### Bioinformatic analysis

The Perseus software platform (26) was used to determine and visualize differential protein abundance between the unventilated control and RM-groups using Welsh’s t-test (two-sided) to account for the variability in group size with p<0.05 considered significant. Protein class composition of significant proteins was assigned using the PANTHER tool (27). Enriched pathways and functions associated with the proteome data sets were identified using the WebGestalt tool (28). As per previous studies (17, 25), to address ovine database limitation in analysis, software homology of sheep to human proteins was assessed by NCBI Basic Local Alignment Search Tool (BLAST); 93% of proteins exhibited homology ≥75%.

#### Physiological and functional parameters

Physiological and functional parameters were recorded at three time points; immediately following the end of the lung recruitment strategy (approximately three minutes post-birth, matched in the No-RM group), and at five minutes and ninety minutes post-birth. Shapiro-Wilk normality testing confirmed the normality of data distribution, with results represented as median and interquartile range. Significant differences between gestation groups were sought using parametric and non-parametric ANOVA as appropriate (one-way ANOVA or Kruskall-Wallis ANOVA, respectively) with post hoc testing to identify intergroup differences as necessary (Holm-Šídák’s or Dunn’s multiple comparison test, respectively). All statistical analysis was performed with GraphPad PRISM 9.1.2 (GraphPad Software, SanDiego, CA) and p<0.05 considered significant. Unless otherwise stated, p values refer to ANOVA.

#### Assessing proteome and physiological phenotypes that differed between the RM groups

To assess if the three treatment groups exhibited differences in the compositional proteome and physiological measures, we used sparse Partial Least Squares (sPLS) regression (29), a multivariate methodology which relates (integrates) two data matrices X (e.g. protein abundance) and Y (e.g. physiological and functional measures), and performs simultaneous variable selection in the two data sets (set at 200 proteins, 20 physiological/functional measures) in the R package MixOmics (version 3.1.1).

### Study Approval

All studies were approved by the Murdoch Children’s Research Institute Animal Ethics Committee in accordance with National Health and Medical Research Council Guidelines.

## AUTHOR CONTRIBUTIONS

P.P-F. Study conception and design, acquisition of data, analysis and interpretation of data, drafting manuscript, statistical analysis. KF. Critical revision of the manuscript for important intellectual content. K.E.McC., R.O., and EP. Acquisition, analysis and interpretation of clinical data, critical revision of the manuscript for important intellectual content. S.B. statistical analysis. D.G.T. Study conception and design, acquisition, analysis and interpretation of clinical data, critical revision of the manuscript for important intellectual content, obtained funding, study supervision. P.P-F. wrote the first draft with assistance from D.G.T. and K.F. All authors approved the final version.

## ACKNOWLEDGEMENTS

The authors thank Mr. Magdy Sourial and Ms. Rebecca Sutton for their assistance with management of the preterm lamb model. This study was supported by a National Health and Medical Research Council Project Grant (1009287) and the Victorian Government Operational Infrastructure Support Program (Melbourne, Victoria, Australia), a National Health and Medical Research Council Clinical Career Development Fellowship (1053889; D.G.T.), and a National Health and Medical Research Council Program Grant (606789). Chiesi Farmaceutici S.p.A. provided the Curosurf used in this study as part of an unrestricted grant (D.G.T.) at the Murdoch Children’s Research Institute.

