## Supplemental Figures and Tables for "Respiratory strategy at birth initiates distinct lung injury phenotypes in the preterm lamb lung"

**SUPPLEMENTARY FIGURE**

**Supplementary Figure 1: Biobank assessment and proteomic sample filtering workflow.**

Abbreviations: dep; gravity dependent, non-dep; non-dependent

Lambs identified in biobank based on aeration streatgy; UVC= 7, No-RM = 19, SI = 26, DynPEEP = 26

UVC= 7, No-RM = 12, SI = 22, DynPEEP = 19

UVC = 7, No-RM = 11 (non-dep lung samples), 9 (dep lung), SI = 22 (non-dep lung), 21 (dep lung), DynPEEP = 16 (non-dep lung), 15 (dep lung)

Poor quality samples identified and removed during Perseus anlaysis

UVC = 6 (non-dep and dep lung samples), No RM = 9 (non-dep and dep lung samples), SI = 19 (non-dep lung), 21 (dep lung), DynPEEP = 15 (non-dep and dep lung samples),

Lung samples removed due to low pepide abundance prior to TMT labelling

Lambs removed from protein extraction due to incomplete functional dataset or ewe illness

**SUPPLEMENTARY TABLES**

**Supplementary Table 1: Clinical characteristics of study population at birth.**

|  | **UVC** | **No-RM** | **SI** | **DynPEEP** | **p value** |
| --- | --- | --- | --- | --- | --- |
| *N* | 7 | 11 | 20 | 19 | N/A |
| Gestational age (days) | 125 (124, 126) | 125 (125, 127) | 126 (124, 127) | 125 (124, 127) | 0.90 |
| Birth weight (kg) | 3.68 (3.32, 4.39) | 3.37 (3.01, 3.58) | 3.38 (3.01, 3.53) | 3.60 (3.21, 3.70) | 0.13 |
| Plurality (S:T) | 0:7 | 2:9 | 7:13 | 2:17 | 0.12 |
| Lung fluid drained (ml/kg) | N/A | 20.8 (15.7, 22.9) | 14.4 (12.3, 18.8) | 16.1 (13.3, 20.6) | 0.16 |
| PaO_2_ (mmHg) | N/A | 25.2 (22.3, 28.5) | 26.5 (23.2, 30.6) | 27.6 (25.8, 29.9) | 0.25 |
| PaCO_2_ (mmHG) | N/A | 47.1 (39.7, 53.0) | 51.9 (45.8, 56.9) | 50.6 (44.1, 53.5) | 0.30 |
| pH | N/A | 7.32 (7.30, 7.40) | 7.35 (7.31, 7.38) | 7.31 (7.30, 7.32)* | 0.035 |
| HCO_3_ (mmol/L) | N/A | 22.8 (22.0, 23.5) | 25.6 (24.0, 26.6)†† | 23.0 (22.4, 23.2)*** | 0.0001 |

Abbreviations: PaO_2_; Fetal partial pressure of O_2_, PaCO2; Fetal partial pressure of CO_2_, pH; Fetal arterial pH, HCO_3_; Fetal arterial bicarbonate

All data median (interquartile range).*p < 0.05 SI *vs.* DynPEEP, ***p < 0.001 SI *vs.* DynPEEP, ††p < 0.01 No-RM *vs.* SI. Gestation age, birth weight, lung fluid drained, pH, bicarbonate; Kruskal-Wallis test. PaO_2_, PaCO_2_; ordinary one-way ANOVA. Plurality; Chi-square contingency test. *Group samples (N) include lambs with either or non-dependent or dependent lung tissue included in analysis*

**Supplementary Table 2: Results from PANTHER protein class composition analysis performed on gravity non-dependent lung samples.**

|  | **No-RM** | | **SI** | | **DynPEEP** | |
| --- | --- | --- | --- | --- | --- | --- |
| **Protein class (PANTHER ID)** | **No. proteins** | **Proportion (%)** | **No. proteins** | **Proportion (%)** | **No. proteins** | **Proportion (%)** |
| Extracellular matrix protein (PC00102) | 6 | 2.9 | 6 | 1.8 | 6 | 2.8 |
| Cytoskeletal protein (PC00085) | 23 | 11.2 | 29 | 8.6 | 20 | 9.3 |
| Transporter (PC00227) | 12 | 5.8 | 22 | 6.5 | 8 | 3.7 |
| Scaffold/adaptor protein (PC00226) | 10 | 4.9 | 19 | 5.6 | 7 | 3.2 |
| Cell adhesion molecule (PC00069) | 1 | 0.5 | 3 | 0.9 | 0 | 0 |
| Nucleic acid metabolism protein (PC00171) | 13 | 6.3 | 27 | 8.0 | 19 | 8.8 |
| Protein-binding activity modulator (PC00095) | 7 | 3.4 | 26 | 7.7 | 17 | 7.9 |
| Calcium-binding protein (PC00060) | 1 | 0.5 | 5 | 1.5 | 4 | 1.9 |
| Gene-specific transcriptional regulator (PC00264) | 4 | 1.9 | 9 | 2.7 | 7 | 3.2 |
| Translational protein (PC00263) | 22 | 10.7 | 40 | 11.8 | 21 | 22.7 |
| Metabolite interconversion enzyme (PC00262) | 46 | 22.3 | 69 | 20.4 | 49 | 22.7 |
| Protein modifying enzyme (PC00260) | 25 | 12.1 | 37 | 10.9 | 23 | 10.7 |
| Chromatin/chromatin-binding, or -regulatory protein (PC00077) | 2 | 1.0 | 5 | 1.5 | 2 | 0.9 |
| Transfer/carrier protein (PC00219) | 4 | 1.9 | 2 | 0.6 | 2 | 0.9 |
| Membrane traffic protein (PC00150) | 19 | 9.2 | 15 | 4.4 | 15 | 6.9 |
| Chaperone (PC00072) | 9 | 4.4 | 19 | 5.6 | 15 | 6.9 |
| Structural protein (PC00211) | 1 | 0.5 | 0 | 0 | 0 | 0 |
| Transmembrane signal receptor (PC00197) | 1 | 0.5 | 3 | 0.9 | 0 | 0 |
| Intercellular signal molecule (PC00207) | 0 | 0 | 1 | 0.3 | 1 | 0.5 |
| Defense/immunity protein (PC00090) | 0 | 0 | 2 | 0.6 | 0 | 0 |

*Grey fill indicates protein classes in which no DEP were identified.*

**Supplementary Table 3: Results from PANTHER protein class composition analysis performed on gravity dependent lung samples.**

|  | **No-RM** | | **SI** | | **DynPEEP** | |
| --- | --- | --- | --- | --- | --- | --- |
| **Protein class (PANTHER ID)** | **No. proteins** | **Proportion (%)** | **No. proteins** | **Proportion (%)** | **No. proteins** | **Proportion (%)** |
| Extracellular matrix protein (PC00102) | 2 | 1.5 | 5 | 3.0 | 10 | 2.9 |
| Cytoskeletal protein (PC00085) | 11 | 8.1 | 22 | 13.3 | 4 | 11.4 |
| Transporter (PC00227) | 4 | 2.9 | 6 | 3.6 | 4 | 11.4 |
| Scaffold/adaptor protein (PC00226) | 7 | 5.2 | 6 | 3.6 | 0 | 0 |
| Cell adhesion molecule (PC00069) | 0 | 0 | 1 | 0.6 | 0 | 0 |
| Nucleic acid metabolism protein (PC00171) | 13 | 9.6 | 15 | 9.1 | 2 | 5.7 |
| Protein-binding activity modulator (PC00095) | 11 | 8.1 | 9 | 5.5 | 0 | 0 |
| Calcium-binding protein (PC00060) | 4 | 2.9 | 2 | 1.2 | 0 | 0 |
| Gene-specific transcriptional regulator (PC00264) | 2 | 1.5 | 5 | 3.0 | 3 | 8.6 |
| Translational protein (PC00263) | 13 | 9.6 | 19 | 11.5 | 3 | 8.6 |
| Metabolite interconversion enzyme (PC00262) | 35 | 25.7 | 37 | 22.4 | 9 | 25.7 |
| Protein modifying enzyme (PC00260) | 13 | 9.6 | 14 | 8.5 | 4 | 11.4 |
| Chromatin/chromatin-binding, or -regulatory protein (PC00077) | 2 | 1.5 | 1 | 0.6 | 0 | 0 |
| Transfer/carrier protein (PC00219) | 2 | 1.5 | 2 | 1.2 | 0 | 0 |
| Membrane traffic protein (PC00150) | 7 | 5.2 | 14 | 8.5 | 4 | 11.4 |
| Chaperone (PC00072) | 5 | 3.7 | 3 | 1.8 | 1 | 2.9 |
| Transmembrane signal receptor (PC00197) | 4 | 2.9 | 3 | 1.8 | 0 | 0 |
| Intercellular signal molecule (PC00207) | 1 | 0.7 | 1 | 0.6 | 0 | 0 |

*Grey fill indicates protein classes in which no DEP were identified.*
